# Functional dissection of neural connectivity in generalized anxiety disorder

**DOI:** 10.1101/2022.01.09.475543

**Authors:** Jonas L. Steinhäuser, Adam R. Teed, Obada Al-Zoubi, René Hurlemann, Gang Chen, Sahib S. Khalsa

## Abstract

Differences in the correlated activity of networked brain regions have been reported in individuals with generalized anxiety disorder (GAD) but an overreliance on the null-hypothesis significance testing (NHST) framework limits the identification and characterization of disorder-relevant relationships. In this preregistered study, we applied a Bayesian statistical framework as well as NHST to the analysis of resting-state fMRI scans from females with GAD and demographically matched healthy comparison females. Eleven *a-priori* hypotheses about functional correlativity (FC) were evaluated using Bayesian (multilevel model) and frequentist (*t*-test) inference. Reduced FC between the ventromedial prefrontal cortex (vmPFC) and the posterior-mid insula (PMI) was confirmed by both statistical approaches. FC between the vmPFC-anterior insula, the amygdala-PMI, and the amygdala-dorsolateral prefrontal cortex (dlPFC) region pairs did not survive multiple comparison correction using the frequentist approach. However, the Bayesian model provided evidence for these region pairs having decreased FC in the GAD group. Leveraging Bayesian modeling, we demonstrate decreased FC of the vmPFC, insula, amygdala, and dlPFC in females with GAD. Exploiting the Bayesian framework revealed FC abnormalities between region pairs excluded by the frequentist analysis, as well as other previously undescribed regions, demonstrating the benefits of applying this statistical approach to resting state FC data.

## Introduction

Generalized anxiety disorder (GAD) is a psychiatric disorder characterized by disproportionate and uncontrollable worry in addition to somatic symptoms including muscle tension, sleep disturbances, fatigue, and difficulty concentrating (American Psychiatric Association, 2013; Barlow, Blanchard, Vermilyea, Vermilyea, & DiNardo, 1986). It is the most common anxiety disorder (Kessler & Wang, 2008), and is associated with substantial functional impairments and economic costs as well as high rates of comorbidity with other psychiatric disorders (Hoge, Ivkovic, & Fricchione, 2012). While the neurobiology of GAD has been investigated extensively (Connor & Davidson, 1998; Stein, 2009), technical advancements in functional neuroimaging in recent decades have afforded insights into abnormalities of regional and network-level neural communication underlying this condition (Fonzo & Etkin, 2017; Holzschneider & Mulert, 2011). Many studies examining brain activity during resting conditions have suggested that specific regions of the brain are organized into functionally dissociable networks that exhibit concurrent activation under varying cognitive conditions, and it is conceivable that functional interdependence between these networks, or nodes within them, is impaired in GAD (Hilbert, Lueken, & Beesdo-Baum, 2014; Kolesar, Bilevicius, Wilson, & Kornelsen, 2019). Among the most frequently described neural networks are the default mode network (DMN, active during the absence of a specific task) (Gusnard, Raichle, & Raichle, 2001; Raichle, 2015), the salience network (SN, responsible for shifting attention to behaviorally relevant internal and external stimuli) (Menon, 2015; Seeley et al., 2007), and the central executive network (CEN, involved in cognitively demanding functions like management of attention) (Bressler & Menon, 2010; Sridharan, Levitin, & Menon, 2008).

Functional connectivity analysis is the most common technique for evaluating neural communication at the network level, which involves assessing temporally dependent co-activation of anatomically separated brain regions during resting conditions (van den Heuvel & Hulshoff Pol, 2010). To reduce causal interpretations of this correlational relationship (Mehler & Kording, 2020), we instead use the term “functional correlativity” (FC). Extant studies on FC in GAD have suggested abnormal relationships between specific brain regions, including the ventromedial prefrontal cortex (vmPFC) (Etkin, Prater, Schatzberg, Menon, & Greicius, 2009; Porta-Casteràs et al., 2020), the insular cortex (Cui et al., 2020; Kolesar et al., 2019; Qiao et al., 2017), the amygdala (Etkin et al., 2009; Hilbert et al., 2014; Kolesar et al., 2019; Li et al., 2016; Roy et al., 2013), and the dorsolateral prefrontal cortex (dlPFC) (Etkin et al., 2009). Additionally, analyses of both task related and resting state functional magnetic resonance imaging (fMRI) data suggest that GAD is characterized by abnormal responses in the dorsal anterior cingulate cortex (dACC) (Blair et al., 2012; Paulesu et al., 2010), the dorsomedial prefrontal cortex (dmPFC) (Mochcovitch, da Rocha Freire, Garcia, & Nardi, 2014), the posterior cingulate cortex (PCC) (Andreescu, Sheu, Tudorascu, Walker, & Aizenstein, 2014), and the temporal pole (TP) (Li et al., 2016). The aforementioned brain regions have been associated with a variety of mental processes including the regulation of emotion (e.g., vmPFC, amygdala) (Hiser & Koenigs, 2018; Phelps, 2006), interoception (e.g., insula) (Craig, 2002; Khalsa et al., 2018), attention (e.g., PCC) (Leech & Sharp, 2014), executive functioning (e.g., dlPFC) (Mansouri, Tanaka, & Buckley, 2009; Petrides, 2000), decision making (e.g., dACC, vmPFC) (Bechara, Tranel, & Damasio, 2000; Bush et al., 2002), working memory (e.g., dlPFC) (Barbey, Koenigs, & Grafman, 2013; Petrides, 2000), processing of mental states (e.g., dmPFC) (Bzdok et al., 2013), and theory of mind (e.g., TP) (Gallagher & Frith, 2003), and accordingly, many of these regions are components of the DM, SN, and CEN.

To date, most studies on resting state FC in GAD have selectively interrogated relationships between subsets of brain regions, often relying purely on the null-hypothesis significance testing (NHST) framework. While frequentist inference requires several assumptions, one of them is particularly problematic in the context of neuroimaging: the conventional mass-univariate analysis unrealistically assumes uniform distribution across spatial units (i.e., voxels, regions). As effects across brain tend to approximately follow a normal distribution, the conventional approach suffers from issues such as information loss, overfitting, and artificial dichotomization (Chen et al., 2021). This underscores the need for an additional, if not alternative, way of looking at the data.

Bayesian inference provides a different statistical viewpoint. It is able to assess evidence in the data both for and against the experimental hypotheses, by allowing the researcher to assess for evidence of invariances as well as differences in a variable of interest (Rouder, Speckman, Sun, Morey, & Iverson, 2009; Wagenmakers et al., 2018). In addition, instead of treating each spatial unit as an isolated entity, as in the conventional mass-univariate analysis, Bayesian multilevel modeling integrates all spatial units into one holistic framework in which all the information is shared and leveraged through partial pooling (Chen et al., 2021). The recent implementation of the multilevel Bayesian modeling matrix-based analysis program (MBA) in AFNI (Cox, 1996) is one such example, which enables researchers to infer the probability of a research hypothesis given the data while overcoming the issue of multiplicity (Chen et al., 2019; Scott & Berger, 2006).

In this preregistered study, we applied a Bayesian statistical framework to the analysis of resting state FC in GAD, with the addition of a frequentist analysis for a conventional comparison. We assessed the FC of brain regions commonly implicated in GAD (vmPFC, dmPFC, dlPFC, dACC, insula, amygdala, PCC, TP) with respect to a set of research hypotheses regarding potential group differences relative to healthy comparisons (HC) (Table 1). Thus in addition to testing hypotheses stemming from the prior frequentist literature on GAD, the application of multilevel Bayesian modeling enabled us to more effectively address issues associated with NHST (such as the problem of multiplicity) and to evaluate observed relationships for convergence (i.e., to functionally “dissect” the data) across analysis approaches.

**Table 1:** *A-priori* hypotheses about differences in functional correlativity between pre-defined regions of interest in generalized anxiety disorder relative to healthy comparisons.

| Region A | Region B | Hypothesis on FC |
| --- | --- | --- |
| Posterior cingulate cortex | Ventromedial prefrontal cortex | Decreased |
| Posterior cingulate cortex | Dorsomedial prefrontal cortex | Decreased |
| Anterior insula | Dorsal anterior cingulate cortex | Decreased |
| Dorsolateral prefrontal cortex | Amygdala | Increased |
| Anterior insula | Ventromedial prefrontal cortex | Decreased |
| Posterior-mid insula | Ventromedial prefrontal cortex | Decreased |
| Amygdala | Dorsal anterior cingulate cortex | Decreased |
| Amygdala | Temporal pole | Increased |
| Amygdala | Ventromedial prefrontal cortex | Increased |
| Amygdala | Anterior insula | Increased |
| Amygdala | Posterior/mid insula | Increased |

Bilateral regions of interest (ROIs) were defined according to the label groupings from the Brainnetome atlas (Fan et al., 2016): Posterior cingulate cortex, ventromedial prefrontal cortex, dorsomedial prefrontal cortex, dorsal anterior cingulate cortex, anterior insula (encompassing the agranular insula in entirety), posterior-mid insula (encompassing the granular and dysgranular insula in entirety), amygdala, dorsolateral prefrontal cortex, temporal pole. The corresponding IDs from the Brainnetome atlas defining each ROI are listed in the materials and methods section.

## Results

### Sample characteristics

Demographic and clinical information of the female study sample are summarized in Table 2. The propensity matching process of study participants (based on body mass index (BMI) and age demographics) resulted in a sample of 29 GAD and 29 HC participants. Three were excluded from further analysis due to excessive motion or signal outliers during their resting scan, resulting in a final analysis sample of 27 GAD and 28 HC participants (for further details on participant selection and recruitment, please refer to the materials and methods section). The participants did not differ significantly on age or BMI, and as expected, the GAD group exhibited higher psychopathology scores on the Patient Health Questionnaire depression module (PHQ-9) (Williams, 2014), Overall Anxiety Severity and Impairment Scale (OASIS) (Campbell-Sills et al., 2009), GAD-7 (Spitzer, Kroenke, Williams, & Löwe, 2006), State-Trait Anxiety Inventory (STAI) (Spielberger, Gorsuch, Lushene, Vagg, & Jacobs, 1983), and Anxiety Sensitivity Index (ASI) (Reiss, Peterson, Gursky, & McNally, 1986) questionnaires (Table 2). The study groups did not differ with respect to average head motion during the resting state scan as indicated by the Wilcoxon signed-rank test (*M_GAD_ =* 0.054, 95% CI [0.044, 0.065]; *M_HC_* = 0.057, 95% CI [0.046, 0.068]; *W* = 393, *p* = 0.809, Δ*M* = 0.003, 95% CI [-0.011, 0.017).

**Table 2:** Demographic and clinical characteristics of study sample.

| | GAD<br>*n = 27 females* | HC<br>*n = 28 females* | $\Delta M$ | 95% CI | *df* | *t* | *p* |
| --- | --- | --- | --- | --- | --- | --- | --- |
| Age (years) | 26.2 ± 6.5 | 24.2 ± 5.1 | -1.99 | [-5.12, 1.18] | 49.23 | -1.26 | 0.214 |
| BMI (kg/m<sup>2</sup>) | 25.9 ± 4.7 | 24 ± 3.2 | -1.86 | [-4.04, 0.32] | 45.5 | -1.72 | 0.092 |
| PHQ-9 | 11.5 ± 5 | 0.7 ± 1.1 | -10.8 | [-12.82, -8.78] | 28.37 | -10.96 | < 0.001 |
| OASIS | 11 ± 2.2 | 1.1 ± 1.6 | -9.82 | [-10.87, -8.78] | 46.97 | -18.9 | < 0.001 |
| GAD-7 | 13.7 ± 3.4 | 1 ± 1.5 | -12.74 | [-14.19, -11.29] | 35.93 | -17.78 | < 0.001 |
| STAI-State | 44.1 ± 9.3 | 24.7 ± 5.6 | -19.37 | [-23.59, -15.15] | 42.48 | -9.26 | < 0.001 |
| STAI-Trait | 58 ± 6.7 | 28.2 ± 6.9 | -29.81 | [-33.51, -26.12] | 51.93 | -16.17 | < 0.001 |
| ASI-Total | 28 ± 14.6 | 7.2 ± 4 | -20.71 | [-26.67, -14.76] | 29.74 | -7.1 | < 0.001 |
| ASI-Physical | 6.3 ± 4.8 | 1.4 ± 1.7 | -4.87 | [-6.85, -2.88] | 31.98 | -4.99 | < 0.001 |
| ASI-Cognitive | 6.5 ± 6.6 | 0.9 ± 1.6 | -5.63 | [-8.29, -2.96] | 28.85 | -4.32 | < 0.001 |
| ASI-Social | 15.1 ± 6.7 | 4.9 ± 2.6 | -10.22 | [-13.01, -7.43] | 33.55 | -7.44 | < 0.001 |

All values presented as means ± standard deviation. Differences between group means ( $\Delta M$ ) and corresponding 95% uncertainty intervals (CI) are reported. Welch's independent samples *t*-tests were conducted to assess differences between the groups. Data for the STAI was missing from one HC participant. *GAD* generalized anxiety disorder, *HC* healthy comparison, *df* degrees of freedom, *BMI* body mass index, *PHQ-9* Patient Health Questionnaire-9, *OASIS* Overall Anxiety Severity and Impairment Scale, *GAD-7* 7-item generalized anxiety scale, *STAI* State-Trait Anxiety Inventory, *ASI* Anxiety Sensitivity Index

### Resting-state fMRI analysis

#### Bayesian multilevel modeling

The results obtained from administering the Bayesian multilevel model (BML) (Chen et al., 2019) to our data identified several region pairs with strong evidence for the group difference being greater (or less) than 0, most notably the vmPFC-PMI region pair (*P_+_* = 0.98). The BML indicated strong evidence for a group difference for some of the region pairs that were also tested in the NHST model (i.e., dlPFC-amygdala, *P*_+_ = 0.96; vmPFC-AI, *P*_+_ = 0.95; PMI-amygdala, *P*_+_ = 0.95).

Interestingly, the BML analysis identified other region pairs to have abnormal FC that were not hypothesized *a-priori* and were therefore not examined in the NHST analysis. These included the dlPFC-PMI, the vmPFC-dACC, the dmPFC-PMI, the dlPFC-dACC, the TP-PMI, and the vmPFC-dmPFC region pairs, all of which indicated high probabilities for a group difference. The complete results of the BML for all region pairs are illustrated in Figure 1.

**Figure 1:**
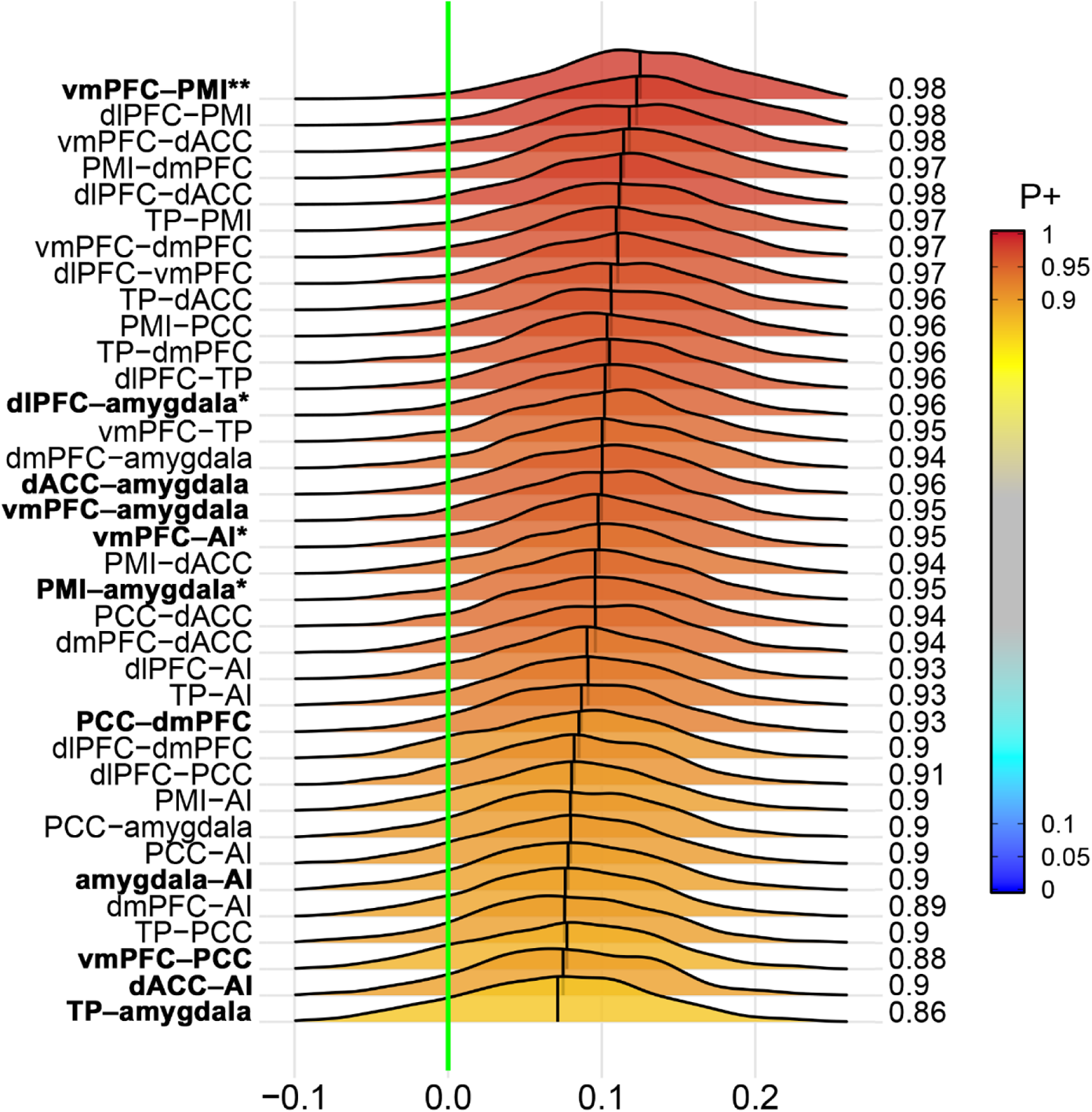
Posterior density distributions of the difference in region-pair effect magnitudes between the two study groups as revealed by the Bayesian multilevel analysis. The value at the right end of each curve indicates the posterior probability *P_+_* for the group difference of the effect being greater than 0 (indicated by the vertical green line). The posterior probability *P_+_* is additionally color-coded in the plane under each posterior density. The vertical black line in each distribution represents the mean effect difference between the two groups for each region pair. Bold font indicates region pairs included in the NHST analysis, with single asterisks indicating significance in the NHST analysis before, and two asterisks for after, Bonferroni-correction for multiple comparisons. *vmPFC* ventromedial prefrontal cortex, *PMI* posterior-mid insula, *dlPFC* dorsolateral prefrontal cortex, *dACC* dorsal anterior cingulate cortex, *dmPFC* dorsomedial prefrontal cortex, *TP* temporal pole, *PCC* posterior cingulate cortex, *AI* anterior insula

The finding of the vmPFC-PMI region pair showing strong evidence for a group difference in the BML was reinforced by regional effect estimates for the group comparisons: Both the vmPFC (*P_+_* = 0.972) and the PMI (*P_+_* = 0.972) showed the highest posterior probabilities of observing a region effect in the HC minus GAD contrast of all areas included in the BML. The complete list of region effects and their respective probabilities are visualized in Figure 2.

**Figure 2:**
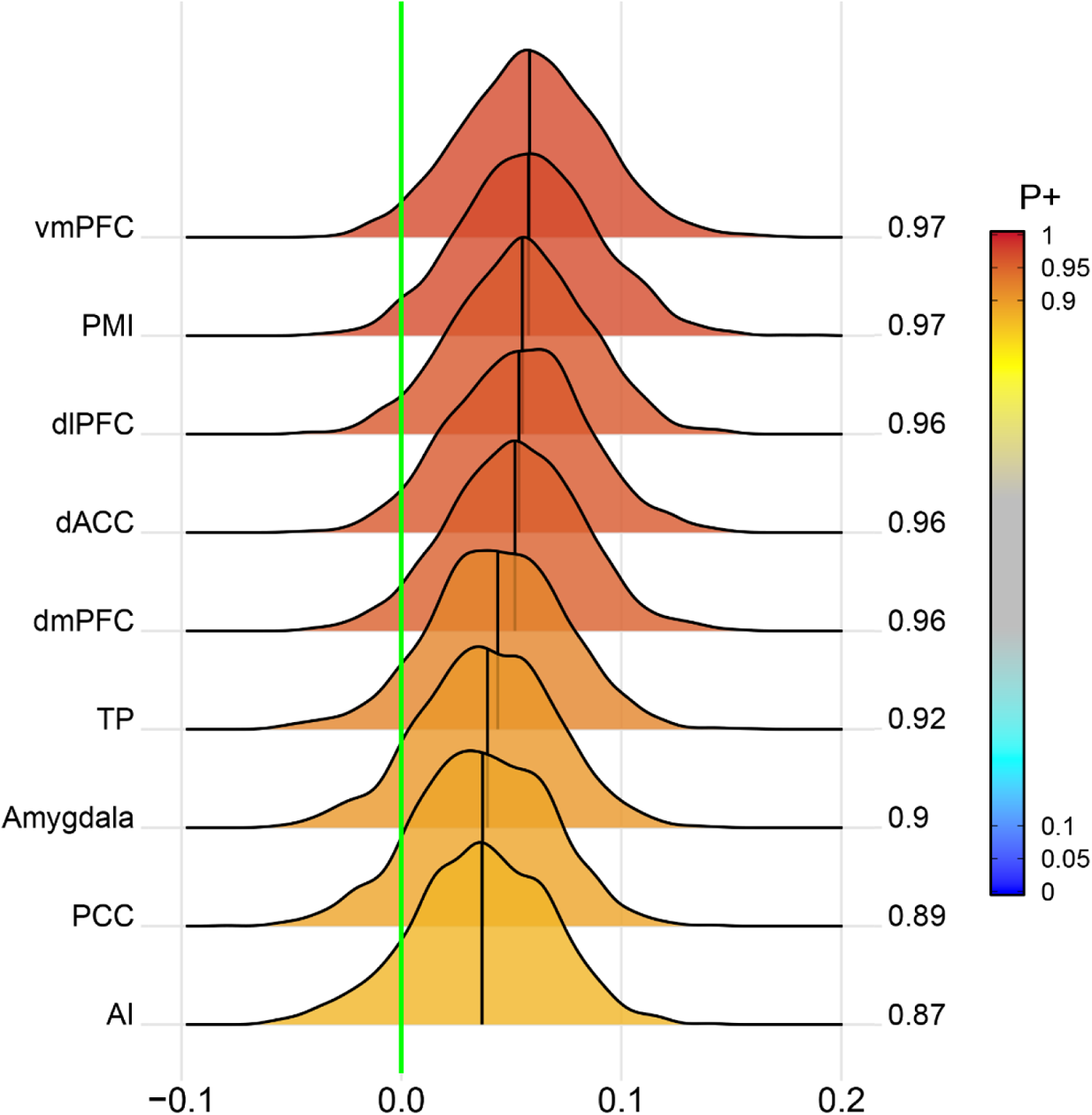
Bayesian multilevel analysis reveals the vmPFC and PMI to have strong evidence for a difference in region effects between the study groups. Posterior density distributions of the difference in region effects in the HC minus GAD contrast of the Bayesian multilevel model. The value at the right end of each curve indicates the posterior probability *P_+_* for the group difference of the effect being greater than 0 (indicated by the vertical green line). The posterior probability *P_+_* is additionally color-coded in the plane under each posterior density. The vertical black line in each distribution represents the mean difference in region effects (as Fishers’s *z*-score) between the two groups for each region in the model. *vmPFC* ventromedial prefrontal cortex, *PMI* posterior-mid insula, *dlPFC* dorsolateral prefrontal cortex, *dACC* dorsal anterior cingulate cortex, *dmPFC* dorsomedial prefrontal cortex, *TP* temporal pole, *PCC* posterior cingulate cortex, *AI* anterior insula

#### Mass-univariate analysis

Results from the conventional mass-univariate analysis revealed that participants in the GAD group had significantly lower FC compared to HCs between the vmPFC and posterior-mid insula (PMI) (*M_GAD_* = 0.29, *SE_GAD_* = 0.05, *M_HC_* = 0.47, *SE_HC_* = 0.04, *t*(48.86) = 2.94, *p* = 0.005), the vmPFC and anterior insula (AI) (*M_GAD_* = 0.41, *SE_GAD_* = 0.04, *M_HC_* = 0.52, *SE_HC_* = 0.03, *t*(52.50) = 2.18, *p* = 0.034), the amygdala and PMI (*M_GAD_* = 0.33, *SE_GAD_* = 0.04, *M_HC_* = 0.43, *SE_HC_* = 0.03, *t*(49.60) = 2.26, *p* = 0.029), and the amygdala and dlPFC (*M_GAD_* = 0.20, *SE_GAD_* = 0.04, *M_HC_* = 0.30, *SE_HC_* = 0.03, *t*(51.52) = 2.16, *p* = 0.036) region pairs. However, after Bonferroni-correction of all hypotheses tested, only the vmPFC and PMI result remained significant (*t*(48.86) = 2.94, *p_adj_* = 0.02) (Figure 3). This finding is in line with the top result from the BML, that convergently identified the vmPFC-PMI region pair to have the highest probability for a group difference. Detailed results for all 11 hypotheses tested can be found in Table 3.

**Figure 3:**
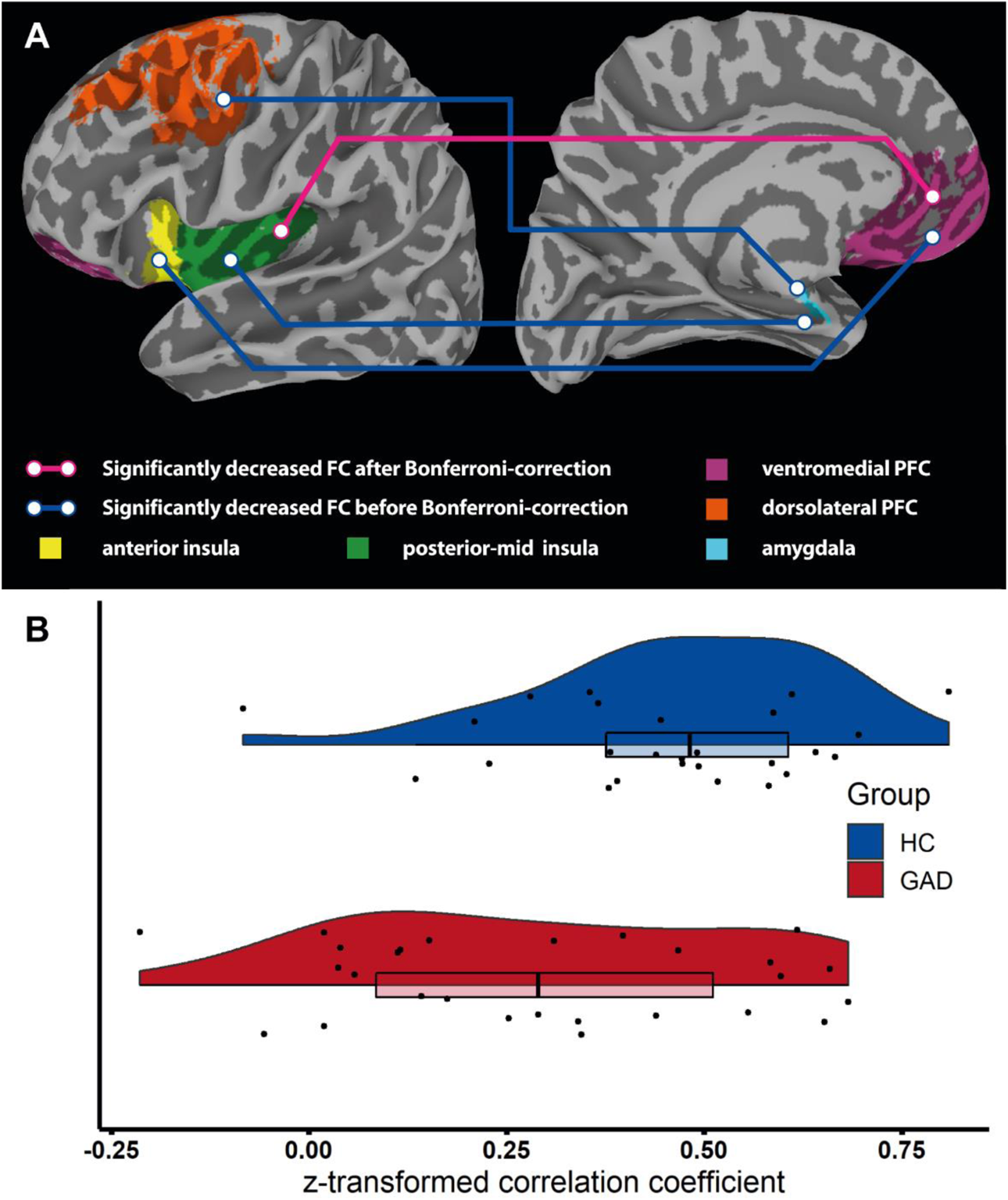
Differences in resting-state FC between GAD and HC revealed by the frequentist analysis. A) The pink line indicates a significantly decreased FC between the PMI and vmPFC in GAD participants after multiple comparison correction using the Bonferroni method. Blue lines indicate differences in FC between the PMI and amygdala, AI and vmPFC, and dlPFC and vmPFC that did not remain statistically significant after Bonferroni correction. Each brain region, indicated by different colors, reflects the selected labels drawn from the Brainnetome atlas. B) Raincloud plots of *Fisher r-to-z* transformed correlation coefficients between the PMI and the vmPFC BOLD-signal time series. *FC* functional correlativity, *GAD* generalized anxiety disorder, *HC* healthy comparison, *vmPFC* ventromedial prefrontal cortex, *PMI* posterior-mid insula, *AI* anterior insula, *dlPFC* dorsolateral prefrontal cortex

**Table 3:**
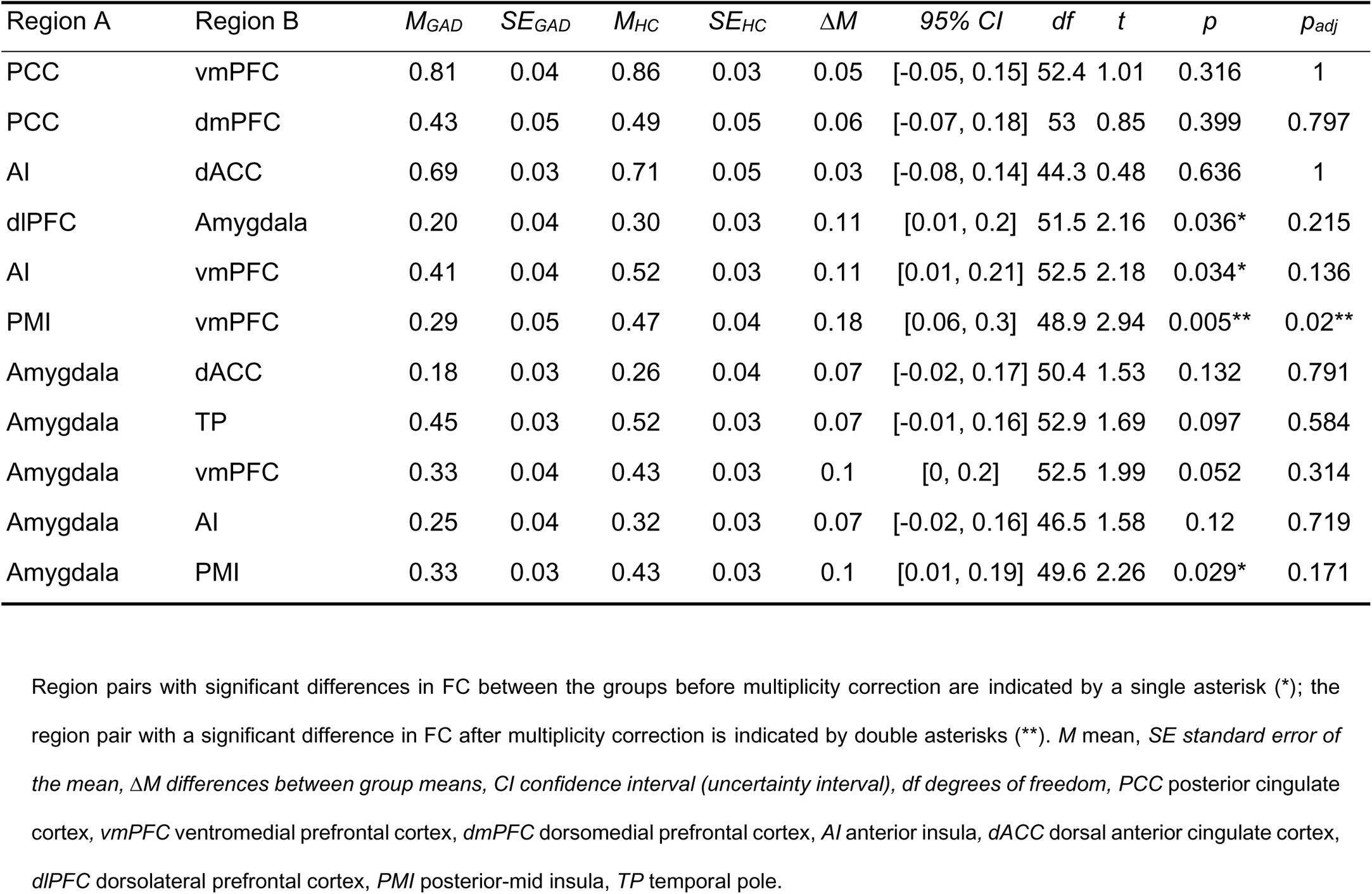
Results of the frequentist analysis of functional correlativity between selected region pairs.

### Exploring the relationship between ventromedial prefrontal cortex - posterior-mid insula correlativity and symptom scores

To investigate whether the reduced FC between the vmPFC and PMI, observed in both frequentist and Bayesian analyses, was related to greater psychopathology we calculated Pearson’s r between vmPFC-PMI z-scores and clinical scores assessed by the following validated questionnaires: The GAD-7, the PHQ-9, the ASI, the STAI, and the OASIS. While the vmPFC-PMI z-score – ASI Total score correlation was statistically significant initially in the GAD group, this relationship did not survive correction for multiplicity. All other correlations were non-significant even before correction for multiple comparisons (Table 4).

**Table 4.**
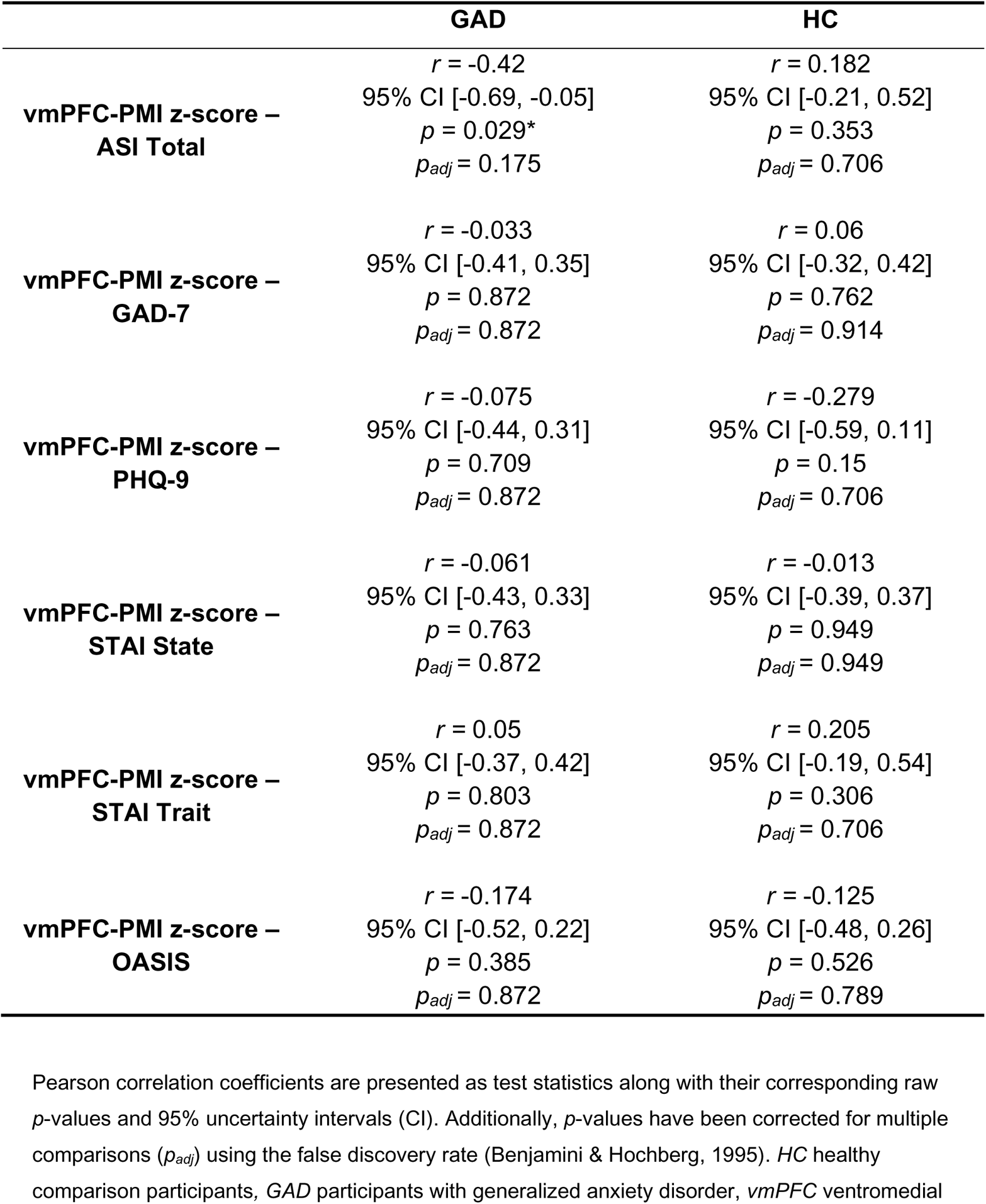
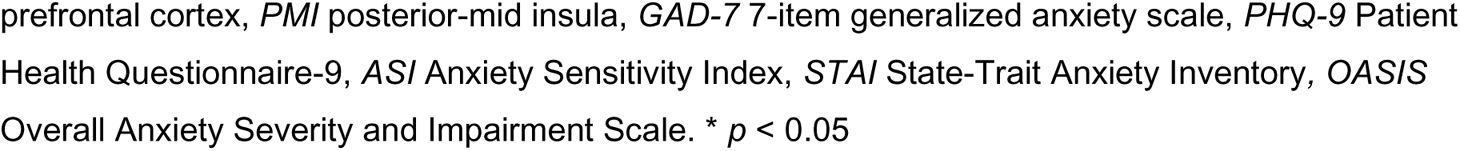
Pearson’s correlation between vmPFC-PMI functional correlativity and clinical variables.

|  | **GAD** | **HC** |
| --- | --- | --- |
| **vmPFC-PMI z-score – ASI Total** | $r = -0.42$<br>95% CI [-0.69, -0.05]<br>$p = 0.029^*$<br>$p_{adj} = 0.175$ | $r = 0.182$<br>95% CI [-0.21, 0.52]<br>$p = 0.353$<br>$p_{adj} = 0.706$ |
| **vmPFC-PMI z-score – GAD-7** | $r = -0.033$<br>95% CI [-0.41, 0.35]<br>$p = 0.872$<br>$p_{adj} = 0.872$ | $r = 0.06$<br>95% CI [-0.32, 0.42]<br>$p = 0.762$<br>$p_{adj} = 0.914$ |
| **vmPFC-PMI z-score – PHQ-9** | $r = -0.075$<br>95% CI [-0.44, 0.31]<br>$p = 0.709$<br>$p_{adj} = 0.872$ | $r = -0.279$<br>95% CI [-0.59, 0.11]<br>$p = 0.15$<br>$p_{adj} = 0.706$ |
| **vmPFC-PMI z-score – STAI State** | $r = -0.061$<br>95% CI [-0.43, 0.33]<br>$p = 0.763$<br>$p_{adj} = 0.872$ | $r = -0.013$<br>95% CI [-0.39, 0.37]<br>$p = 0.949$<br>$p_{adj} = 0.949$ |
| **vmPFC-PMI z-score – STAI Trait** | $r = 0.05$<br>95% CI [-0.37, 0.42]<br>$p = 0.803$<br>$p_{adj} = 0.872$ | $r = 0.205$<br>95% CI [-0.19, 0.54]<br>$p = 0.306$<br>$p_{adj} = 0.706$ |
| **vmPFC-PMI z-score – OASIS** | $r = -0.174$<br>95% CI [-0.52, 0.22]<br>$p = 0.385$<br>$p_{adj} = 0.872$ | $r = -0.125$<br>95% CI [-0.48, 0.26]<br>$p = 0.526$<br>$p_{adj} = 0.789$ |

Pearson correlation coefficients are presented as test statistics along with their corresponding raw $p$ -values and 95% uncertainty intervals (CI). Additionally, $p$ -values have been corrected for multiple comparisons ( $p_{adj}$ ) using the false discovery rate (Benjamini & Hochberg, 1995). *HC* healthy comparison participants, *GAD* participants with generalized anxiety disorder, *vmPFC* ventromedial
prefrontal cortex, *PMI* posterior-mid insula, *GAD-7* 7-item generalized anxiety scale, *PHQ-9* Patient Health Questionnaire-9, *ASI* Anxiety Sensitivity Index, *STAI* State-Trait Anxiety Inventory, *OASIS* Overall Anxiety Severity and Impairment Scale. \* $p < 0.05$

## Discussion

In this preregistered study, we examined FC in females with GAD relative to matched HCs to test a set of *a-priori* hypotheses using dual statistical frameworks: Bayesian multilevel modeling and NHST. Converging results from both analyses confirmed diminished FC between the PMI and the vmPFC in the GAD vs. HC groups. FC between these regions was significantly associated with one clinically relevant trait measure, anxiety sensitivity, in the GAD group, however this relationship did not survive multiple comparison corrections.

While the frequentist analysis in convergence with the BML identified the vmPFC and PMI region pair to exhibit decreased functional coupling in our analysis, the NHST framework as a whole has faced growing criticism (Greenland et al., 2016; Nickerson, 2000; Wasserstein & Lazar, 2016). Conceptually, NHST results are often misinterpreted (or misunderstood) given that they assess only whether the experimental result observed (e.g., that two groups differ on a variable of interest) is too unlikely to maintain an assumption that the null hypothesis is true (Chen et al., 2019; Nickerson, 2000). This assumption precludes the ability to distinguish between results that provide evidence of the absence of an effect from null results yielded merely from a lack in statistical power. Additionally, using the same data to test multiple hypotheses in the NHST framework arbitrarily inflates the type I error (Tukey, 1991), resulting in a variety of methods to adjust for this problem of multiplicity (Bender & Lange, 2001; Curran-Everett, 2000). Examining the data using Bayesian multilevel model overcomes these challenges while further providing evidence for abnormal FC among region pairs that were not hypothesized *a-priori* and were therefore not tested in the NHST analysis. This included evidence of abnormal FC among several regions including the dlPFC-PMI, the vmPFC-dACC, the dmPFC-PMI, the dlPFC-dACC, the TP-PMI, and the vmPFC-dmPFC. Thus, the application of both statistical frameworks allowed for a functional dissection of neural connectivity in GAD yielding confirmatory results (most notably vmPFC-PMI) and providing indications of other relationships worth examining further.

Numerous studies have implicated the vmPFC in key aspects of cognitive functioning such as decision making (Bechara et al., 2001, 2000), generation and regulation of emotion (Diekhof, Geier, Falkai, & Gruber, 2011; Hiser & Koenigs, 2018; Winecoff et al., 2013), and fear conditioning (Quirk & Mueller, 2008; Sotres-Bayon, Cain, & LeDoux, 2006). The vmPFC has been previously associated with greater fear generalization in GAD (Cha et al., 2014; T. Greenberg, Carlson, Cha, Hajcak, & Mujica-Parodi, 2013), which fits the clinical picture of excessive worry in individuals with the disorder (Rowa & Antony, 2008). Moreover, abnormal vmPFC functioning has most often been implicated in anxiety disorders in regards to its proposed role of inhibiting amygdala output ((Davidson, 2002; Phelps, Delgado, Nearing, & LeDoux, 2004); but see (Myers-Schulz & Koenigs, 2012)). This seems reasonable considering the widely accepted view of the amygdala as a central hub for fear processing (Davis, 1992; LeDoux, 2003). However, several lines of evidence show the need to distinguish between exteroceptive fear processing, which is most prominently mediated through the amygdala, and interoceptive fear processing, which is most prominently mediated through the insular cortex. For example, studies of individuals with bilateral amygdala lesions have shown a remarkable absence of anxiety or panic in response to exteroceptive fear stimuli (Adolphs & Tranel, 2000; Bechara et al., 1995; Becker et al., 2012; Feinstein, Adolphs, Damasio, & Tranel, 2011) but experienced fear and panic evoked by interoceptive stimuli (Feinstein et al., 2013; Khalsa et al., 2016). Interoception, which encompasses the sensory processing of internal body signals by the nervous system (Craig, 2002; Khalsa et al., 2018), is a process tightly linked to the insular cortex among other regions including the medial prefrontal cortex and amygdala (Berntson & Khalsa, 2021; Khalsa, Rudrauf, Feinstein, & Tranel, 2009). Models of interoceptive processing have suggested a posterior-to-anterior integration of interoceptive signaling within the human insula (Barrett & Simmons, 2015; Craig, 2009; Seth, Suzuki, & Critchley, 2011) that is supported by its pattern of cytoarchitectonic organization with an agranular rostral and dysgranular/granular mid and posterior divisions across humans and primates (Evrard, Logothetis, & Craig, 2014; Ghaziri et al., 2017; Morel, Gallay, Baechler, Wyss, & Gallay, 2013). Studies examining the functional organization of the insula implicate the AI in task maintenance (Dosenbach et al., 2006), attention control (Nelson et al., 2010), emotion (Cauda et al., 2011; Zaki, Davis, & Ochsner, 2012), and predictive processing (Barrett & Simmons, 2015; Seth et al., 2011), which is in line with increased insula activity during emotional processing tasks in individuals with anxiety disorders (Simmons, Strigo, Matthews, Paulus, & Stein, 2006; Stein, Simmons, Feinstein, & Paulus, 2007). Thus, the reduced vmPFC-PMI FC observed in this current study supports the idea that individuals with GAD may have difficulty exercising top-down regulation of emotion due to aberrant processing of signals flowing through an interoceptive hub: the insula. This hypothesis is backed by the vmPFC and PMI having the highest probabilities for a region effect in the HC minus GAD contrast as well as by FC between the vmPFC and the AI being reduced in the GAD group as convergently confirmed by both statistical approaches (although the NHST-result was only statistically significant before correcting for multiplicity).

Other results from the frequentist analysis indicate abnormal FC of the amygdala: though contrary to our hypothesis, we observed decreased, not increased, FC between the amygdala and the PMI. The direction of this finding contrasts with previous reports of an amygdala-insula resting state network in both anxious adults (Baur, Hänggi, Langer, & Jäncke, 2013) and adolescents (Roy et al., 2013) but on the other hand aligns with previous findings of reduced amygdala-insula FC (Etkin et al., 2009). Additionally, FC between the amygdala and the dlPFC was decreased, not increased, in our GAD sample. This was in divergence with our hypothesis which was based on previous literature (Etkin et al., 2009). Decreased FC between the amygdala and the dlPFC, which is a central node in the CEN (Bressler & Menon, 2010; Menon, 2011), could be argued to reflect a dysfunctional management of attention (a key function of the CEN) (Bressler & Menon, 2010) towards threat-related stimuli, which is certainly a clinical key-feature of GAD (MacLeod, Mathews, & Tata, 1986; Mathews & MacLeod, 1985; Mogg & Bradley, 2005). However, the overly general view of the amygdala as the central hub of fear processing is challenged by the absence of amygdala involvement in human fear extinction in a recent meta-analysis (Fullana et al., 2018) and heterogenous amygdala findings across reviews of neuroimaging literature in GAD (Goossen, van der Starre, & van der Heiden, 2019; Hilbert et al., 2014; Mochcovitch et al., 2014). While the results from our cross-sectional study might hint toward the possibility that the role of the amygdala might not be as pivotal to the maintenance of GAD as expected, both amygdala-related findings (i.e., reduced FC for the PMI-amygdala and the dlPFC-amygdala in the GAD group) did not withstand correction for multiplicity and would therefore not be considered statistically significant using the NHST model framework. On the other hand, evaluation of the results from the BML indicated high probabilities for a group difference regarding those region pairs, raising the question whether overly rigorous multiplicity correction might have induced a type I error in the NHST-analysis of those brain regions. Viewing the data from a different, i.e., Bayesian, perspective thus strengthened the validity of the amygdala findings, permitting us to discuss these results and consider their potential implications for GAD.

Bayesian multilevel modeling further allowed us to investigate relationships in GAD that were not hypothesized *a-priori* with minimal risk of information loss. Along with the converging result of decreased vmPFC-PMI FC from both statistical approaches, our analysis identified high probabilities for decreased FC of the PMI with the dlPFC, the dmPFC, and the TP. Decreased functional coupling of the PMI and the dlPFC could be interpreted to reflect abnormal signaling of internal body signals to a key region for executive functions like working memory (Barbey et al., 2013) and attention (Kane & Engle, 2002): aspects of cognition known to be impaired in anxiety (Bar-Haim, Lamy, Pergamin, Bakermans-Kranenburg, & van IJzendoorn, 2007; Vytal, Cornwell, Letkiewicz, Arkin, & Grillon, 2013). The reduced PMI FC between both the dmPFC (a brain area known to be hyperactivated in GAD during emotional processing (Paulesu et al., 2010) and at rest (Wang et al., 2016)), and the TP (an area implicated in social and emotional processing) (Olson, Plotzker, & Ezzyat, 2007; Wong & Gallate, 2012), align well with a proposed model of the insula as an “integral hub” for detecting salient events, and for switching attention to these stimuli in preparation for regulatory (i.e., visceromotor) processing (Menon & Uddin, 2010).

The Bayesian multilevel analysis also revealed diminished FC of the vmPFC-dmPFC region pair in GAD, two key components of the DMN (Raichle et al., 2001). This finding is consistent with previous reports of DMN alterations in GAD (Andreescu et al., 2014; Zhao et al., 2007), albeit diminished FC between the vmPFC and dmPFC has not been reported previously. These regions of the DMN are hypothesized to promote functions like processing of emotion and self-referential cognition (Raichle, 2015), which are impaired in GAD (Turk, Heimberg, Luterek, Mennin, & Fresco, 2005; Watson, Timulak, & Greenberg, 2019). Lastly, the Bayesian analysis revealed reduced FC with the vmPFC and the dlPFC. Given that the vmPFC and the dlPFC are key components, respectively, of the DMN and CEN networks (Menon, 2015; Seeley et al., 2007), these reductions in FC could disrupt the contribution of the dACC to switching between the spontaneous cognition of the DMN (Andrews-Hanna, Reidler, Huang, & Buckner, 2010) and executive functioning of the CEN (Seeley et al., 2007) and may ultimately result in impaired action selection in those with GAD, a function for which the dACC is critical (Rushworth, 2008).

### Limitations

Our focus on females with GAD was based on the fact that females outnumber males and that our sample was drawn from a larger study examining psychiatric disorders predominantly affecting females (e.g., anorexia nervosa and GAD). Future research is needed to establish whether our findings extend to males, i.e., whether sex differences in FC play a role in GAD; such discrepancies have not been reported in previous GAD studies. While our sample size is above the median (N = 23-24) for fMRI studies in recent years (Szucs & Ioannidis, 2020), it remains modest in light of evolving best practices and estimates of the size necessary to ensure replicability. A general limitation of the FC analysis approach employed here is that it cannot determine the directionality (or responsible region) for impaired functional coupling observed within region pairs. Since we tested hypotheses regarding interrelationships of individual brain regions at rest, we cannot make inferences about network-level FC as commonly assessed by seed-based, voxel-wise whole-brain analyses. Analysis of resting state fMRI data has received growing criticism regarding potential confounds including, most recently, the discovery of “resting state physiological networks” (i.e., physiologically driven FC resembling previously reported neural networks) (J. E. Chen et al., 2020). We addressed this concern by recording and regressing out signals attributed to respiration or cardiac pulsatility (Glover, Li, & Ress, 2000). A recent paper also identified substantial variability of fMRI results across many teams analyzing the same data set (Botvinik-Nezer et al., 2020), underlining the need for standardized fMRI analysis measures. To improve the reproducibility of our findings, we followed several of the recommended steps including 1) applying different statistical approaches to the data yielding largely converging results, 2) pre-registering our hypotheses and statistical approaches before analyzing any study data, 3) reporting results of all analyses conducted, even if they did not reach statistical significance, and 4) publicly sharing the code of our data processing pipeline and statistical analysis (J. L. Steinhäuser, 2021).

## Conclusion

We leveraged the strengths of the Bayesian inference framework to convergently identify reduced FC between the vmPFC and the PMI in GAD. The Bayesian framework allowed us to identify FC abnormalities between region pairs excluded by the frequentist analysis and other previously undescribed regions, emphasizing the utility of this approach for probing the pathophysiological basis of psychiatric disorders. Future fMRI studies of resting state FC may benefit from a similar approach.

## Materials and methods

The study hypotheses and data analysis plan were registered (J. Steinhäuser, Teed, & Khalsa, 2020) on the Open Science Framework (Foster & Deardorff, 2017) before any of the study data was accessed or processed. All study data and analysis scripts are available online (J. L. Steinhäuser, 2021).

### Participants

Females with GAD and female HCs were recruited for this study from the Tulsa metropolitan area via advertisement in newspaper, radio, and social media outlets as well as via outpatient referral from the Laureate Psychiatric Clinic and Hospital. We report data that was collected as part of a larger, ongoing fMRI study that included an interoceptive perturbation task (isoproterenol infusion) performed after collection of the resting data presented here (Teed et al., 2020). Since this large-scale study focuses on psychiatric disorders that predominantly occur in females, the sample base for this investigation was also female-only. Further details on the aforementioned study can be found on the ClinicalTrials.gov registration NCT02615119 at the U.S. National Library of Medicine (Khalsa, 2015). The selection of participants is visualized in a CONSORT diagram (Figure 4). The diagnosis of GAD was verified according to DSM-5 (American Psychiatric Association, 2013) criteria by an experienced clinician administering the M.I.N.I. neuropsychiatric interview (Sheehan et al., 1998). Additional GAD inclusion criteria were a currently elevated level of anxiety as evidenced by a GAD-7 score greater than 10 out of 21 or an OASIS score greater than 7 out of 20. All participants were administered the PHQ-9, the GAD-7 questionnaire, the OASIS, the STAI, and the ASI. For the GAD group, selected psychotropic agents (e.g., serotonergic/noradrenergic) were allowed so long as they were stably medicated (no changes within four weeks). Pro re nata (PRN) medications were not a criterion for exclusion so long as participants were able to abstain from their use for at least two days prior to testing. Any history of a psychotic disorder or bipolar disorder led to exclusion from this study. Because of the cardiovascular implications of the pharmacological task employed as part of the larger study, participants previously diagnosed with cardiac or respiratory diseases were excluded, as well as those with comorbid panic disorder. Individuals in the GAD group had the following psychiatric comorbidities: 11/27 Major depressive disorder (MDD), 9/27 MDD and social anxiety disorder (SAD), 1/27 SAD, 1/27 MDD and post-traumatic stress disorder (PTSD), 1/27 MDD and alcohol use disorder (mild), 1/27 SAD and obsessive-compulsive disorder, 1/27 MDD, SAD, and agoraphobia, 1/27 MDD, SAD, and PTSD. Individuals in the GAD group reported taking the following psychoactive medication: 3/27 selective serotonin reuptake inhibitor (SSRI), 2/27 selective norepinephrine reuptake inhibitor (SNRI), 1/27 SSRI/SNRI, and 1/27 medicinal tetrahydrocannabinol. HCs were required to be without any history of psychiatric illness per the M.I.N.I. interview. The HC group was individually matched to the GAD group so that they would not differ significantly on BMI and age due to the known influence of the former on head motion (Ekhtiari, Kuplicki, Yeh, & Paulus, 2019; Van Dijk, Sabuncu, & Buckner, 2012) and the latter on FC (Betzel et al., 2014; Geerligs, Renken, Saliasi, Maurits, & Lorist, 2015). The study was approved by the Western institutional review board and was conducted at the Laureate Institute for Brain Research in accordance with the Declaration of Helsinki. All participants provided written informed consent and received financial compensation for their study involvement.

**Figure 4:**
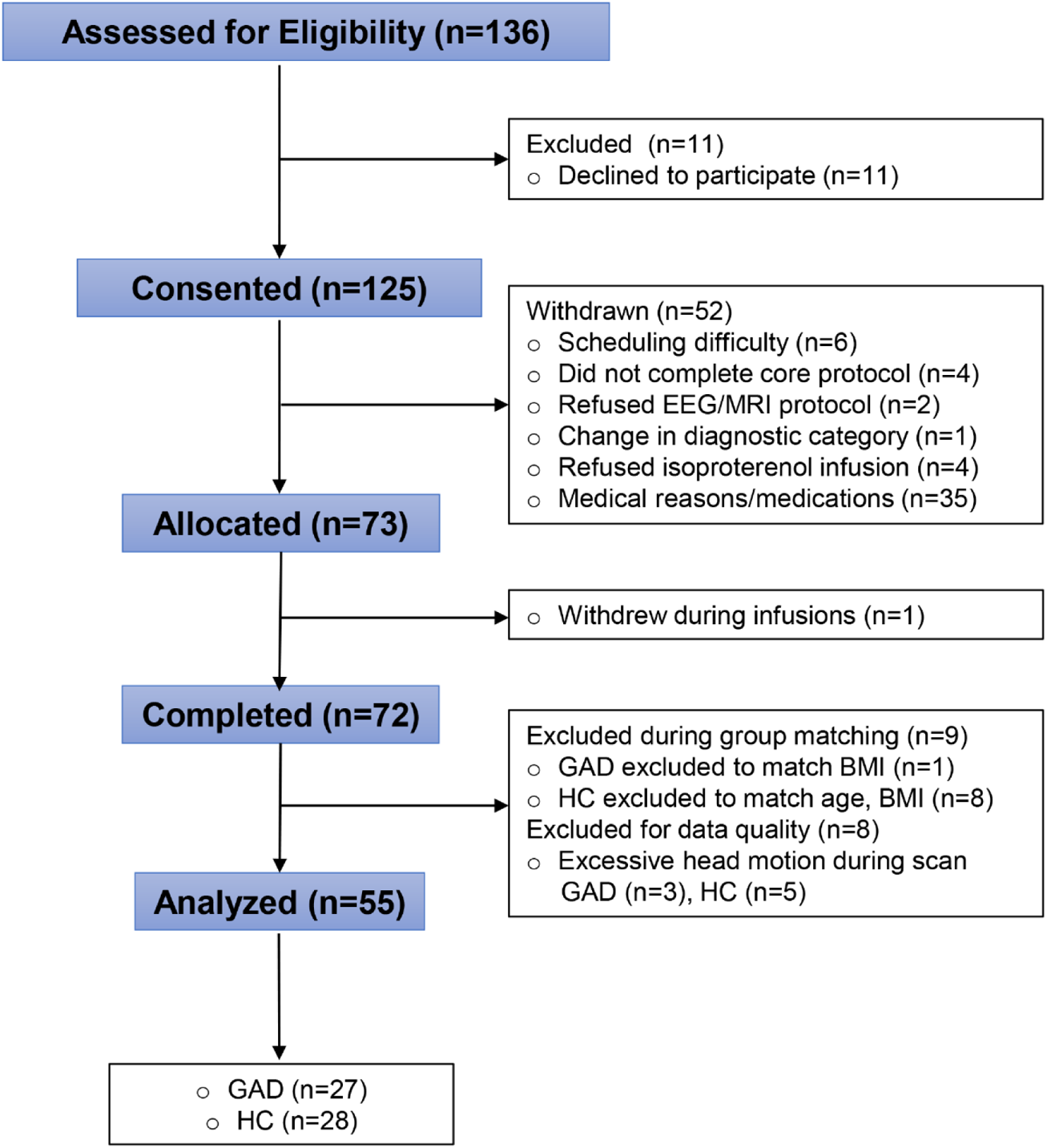
CONSORT diagram of the isoproterenol (ISO) study and its resting state data analyzed for this investigation. Adapted with permission from Teed et al. (2020). *HC* healthy comparison participants, *GAD* participants with generalized anxiety disorder, *ISO* isoproterenol, *rsfMRI* resting state functional magnetic resonance imaging

### Image acquisition

Magnetic resonance images were obtained using two identical full-body 3.0 Tesla MR750 MRI scanners (GE Healthcare, Milwaukee, WI), equipped with an 8-channel receive-only head array coil (GE Healthcare, Milwaukee, WI). First, a T1-weighted image was acquired as an anatomical reference using a magnetization-prepared rapid gradient echo (MPRAGE) sequence with sensitivity encoding (SENSE) (Pruessmann, Weiger, Scheidegger, & Boesiger, 1999) over the duration of 5 minutes and 40 seconds. The sequence-parameters were: FOV = 240×192 mm, matrix = 256×256, 186 axial slices, slice thickness = 0.9 mm, 0.938×0.938×0.9 mm^3^ voxel volume, TR = 5 ms, TE = 2.012 ms, SENSE acceleration factor R = 2, flip angle = 8°, delay time = 1400 ms, inversion time = 725 ms, sampling bandwidth = 31.25 kHz. Next, a resting-state scan was conducted using a single-shot gradient-recalled echo-planar imaging (EPI) sequence with SENSE and the following parameters: TR = 2000 ms, TE = 27 ms, R = 2, FA = 78°, FOV = 240 mm, 39 axial slices with 2.9 mm thickness with no gap, matrix = 96×96. The EPI images were reconstructed into a 128×128 matrix that produced 1.875×1.875×2.9 mm^3^ voxel volume. Prior to the resting-state scan, participants were instructed to remain as still as possible, to keep their eyes open and fixated on a cross presented at the center of the screen, and to “clear your mind and do not think about anything in particular”. During the scan, respiration was recorded using a pneumatic belt placed around the torso. Heart rate was recorded using a photoplethysmograph with an infrared emitter placed under the pad of the participant’s finger.

### Data analysis

#### Preprocessing

Preprocessing of fMRI data was conducted using AFNI 20.0.19 (Cox, 1996, RRID:SCR_005927) and Freesurfer 6.0.0 (Fischl, 2012, RRID:SCR_001847). T1-weighted images were skull stripped and nonlinearly warped to Montreal Neurological Institute (MNI) 152 atlas space. White matter and ventricle masks were acquired to later regress out their signal from the data. The first three images of each participant’s timeseries were removed to ensure an equilibrium of fMRI signal. EPI volume signal was despiked using AFNI’s *3dDespike* program with default parameters. Physiological noise effects (i.e., due to cardiac pulsatility and respiration) (Glover & Lee, 1995; Noll & Schneider, 1994) were regressed out using the RETROICOR method (Glover et al., 2000) implemented in AFNI. Slice timing correction was performed to account for interleaved slice acquisition. Anatomical images and EPI volumes for each participant were aligned to their EPI volume determined to have the minimum outliers according to a *Local Pearson Correlation Signed* cost function in AFNI. Datasets were blurred using a Gaussian kernel with full width half maximum of 4 mm. The time series of each voxel was scaled to a mean of 100, so that values could be interpreted as percentage change from the mean. Subsequently, voxel-values with a percentage increase of ≥ 100% were removed as outliers. Volumes censored due to too much motion or being signal outliers were interpolated using the previous and subsequent volume. Participants displaying excessive motion or signal outliers during their resting scan (i.e., > 30% volumes being censored because of motion or signal outliers) were excluded.

#### Region of interest definition and data extraction

Based on a careful review of the fMRI literature on GAD we assessed FC between a total of nine regions of interest (ROIs): vmPFC, PMI, dlPFC, dACC, dmPFC, TP, amygdala, PCC, and AI. We then formulated a total of 11 *a-priori* hypotheses about FC between the nine pre-defined ROIs for examination (Table 1). To extract the data for each ROI, a mask was created by collapsing over the relevant labels of the Brainnetome atlas (Fan et al., 2016), which provides a probabilistic cytoarchitectonic parcellation of the human brain. The average timeseries for each ROI was then extracted for each participant using AFNI’s 3dROIstats program. The following IDs from the Brainnetome atlas were used to create the ROI analysis mask: PCC: *153, 154, 175, 176, 181, 182*; vmPFC: *41, 42, 45, 46, 47, 48, 49, 50, 187, 188*; dmPFC: *1, 2, 11, 12*; dACC: *179, 180, 183, 184*; AI, encompassing the agranular insula in entirety: *165, 166, 167, 168*; PMI, encompassing the granular/dysgranular insula in entirety: *163, 164, 169, 170, 171, 172, 173, 174*; amygdala: *211, 212, 213, 214*; dlPFC: *3, 4, 15, 16, 17, 18, 23, 24, 25, 26*; TP: *69, 70*. The resulting ROI analysis mask is supplied in MNI-space and AFNI format with this manuscript (J. L. Steinhäuser, 2021).

#### Statistical analyses

Using the timeseries of our nine ROIs we constructed a 9 x 9 correlation matrix for each participant. The relationship between ROIs was assessed using Pearson’s correlation. The resulting sampling distribution of Pearson’s *r* was normalized using the *Fisher r-to-z transformation* and the obtained z-scores were used in all further analyses.

### Bayesian modeling

A Bayesian multilevel model (BML) (Chen et al., 2019) was applied to our data using the MBA program in AFNI, estimating the posterior probability of the effect being greater than 0 (*P_+_*). The BML was also used to explore all other possible region pairs that we did not hypothesize *a-priori* to be aberrant in GAD, and therefore left out of the FC analysis. The BML overcomes limitations of NHST in this context by (a) incorporating the interrelationships between region pairs into one model through partial pooling, (b) addressing the issue of multiplicity in a NHST framework, (c) providing direct evidence for or against the effect of a region pair instead of assuming the null-hypothesis (Chen et al., 2019), (d) estimating the contribution of each individual brain region in the network relative to all other regions as a measurement of “relative importance”, and (e) supporting full result reporting and treating statistical evidence as a continuum instead of arbitrary dichotomization.

### Mass univariate analysis

Welch’s independent samples *t*-tests were used to test the null hypothesis that there was no difference in FC-scores between the two groups. To prevent the inflation of Type I errors (i.e., the problem of multiplicity) the results were Bonferroni corrected. To decrease the likelihood of committing Type II errors, only the region pairs hypothesized to be aberrant in GAD (Table 1) were tested. Since some brain regions were included in more hypotheses than others, their data was used in multiple comparisons. The amygdala was included in six, the vmPFC in four, the AI in three, the PMI, PCC, and dACC in two, and the dmPFC, dlPFC, and TP in one comparison(s). Consequently, the significance level of the test results for each region pair were corrected based on how many comparisons the regions were included in. Finally, exploratory relationships between FC and symptom scores were examined using Pearson’s correlation.

## Acknowledgements

The authors wish to acknowledge the support of Valerie Upshaw MSN, APRN-CNP in gathering participants information and Rayus Kuplicki PhD for help with MRI data management.

## Financial support

Funding statement: This work was supported by National Institute of General Medical Sciences (NIGMS) Center Grant P20GM121312 (S.S.K.), National Institute of Mental Health Grant K23MH112949 (S.S.K.), The William K. Warren Foundation (S.S.K.) and the German Federal Ministry of Education and Research by providing J.S. with a scholarship for his collaboration with the Laureate Institute for Brain Research. The views expressed in this article are those of the authors and do not necessarily reflect the position or policy of the National Institutes of Health.

## Competing interests

The authors declare no competing interests.

## Ethical standards

The authors assert that all procedures contributing to this work comply with the ethical standards of the relevant national and institutional committees on human experimentation and with the Helsinki Declaration of 1975, as revised in 2008.

## Notes

https://github.com/Jonas-Ste/GAD_MBA_FC

### Competing Interest Statement

The authors have declared no competing interest.

